# The extracellular matrix gene *mec-9* regulates *C. elegans* sensory cilia

**DOI:** 10.64898/2026.03.13.711665

**Authors:** Katherine C. Jacobs, Deanna M. De Vore, Karla M. Knobel, Jonathon D. Walsh, Alakananda Das, Leah M. Dobossy, Inna A. Nikonorova, Ken C. Q. Nguyen, Miriam B. Goodman, David H. Hall, Maureen M. Barr

**Affiliations:** Department of Genetics and Human Genetics Institute of New Jersey, Rutgers, The State University of New Jersey, Piscataway, NJ, USA; Diane C. Lobosco STEM Academy, Passaic County Vocational Schools, Wayne, NJ, USA; Waisman Center, University of Wisconsin-Madison, Madison, WI, USA; Department of Molecular and Cellular Physiology, Stanford University School of Medicine, Stanford, CA, USA; Center for C. elegans Anatomy, Albert Einstein College of Medicine, Bronx, NY, USA

## Abstract

Cilia are critical sensory organelles that project from the cell surface into the tissue environment, where they are surrounded by extracellular matrix (ECM). Abnormal ECM and fibrosis are two hallmarks of ciliopathies, yet the relationship between cilia and ECM is not well understood. Using the sense organs of *C. elegans* as a model, we found that a neomorphic mutation in the ECM gene *mec-9* impacts sensory cilia function, ciliary protein localization, microtubule ultrastructure, and shedding of ciliary extracellular vesicles (EVs). We show that *mec-9* is not expressed in EV releasing neurons, but rather by companion neurons in the sense organs, and may act cell non-autonomously. Our studies reveal pleiotropic roles for *mec-9* in the *C. elegans* ciliated nervous system and provide an *in vivo* model to study the relationship between cilia and ECM.

## Introduction

Sensory cilia are microtubule-based membrane protrusions enriched for receptors, channels, and signaling components that many eukaryotic cells use to sense and respond to their environment (Hilgendorf et al., 2024). Cilia also transmit signals by releasing extracellular vesicles (EVs) (Wang et al., 2024a). The importance of cilia in human health and disease is demonstrated by a class of human diseases known as ciliopathies, in which cilia formation or function is disrupted (Hilgendorf et al., 2024). Cilia project into the tissue environment where they are surrounded by extracellular matrix (ECM), a secreted substance composed of proteins, lipids, and proteoglycans that supports, protects, and sends signals to cells (Frantz et al., 2010; Naba, 2024). Ciliary signaling can influence ECM production (Sieckmann et al., 2025; Stevenson, 2023) and abnormal ECM remodeling and fibrosis occur in many ciliopathies (Alzarka et al., 2025), thus understanding the relationship between cilia and ECM is critically important.

ECM components and regulators have been shown to regulate ciliogenesis (Failler et al., 2021) and cilia structural organization (Yu et al., 2016). For example, in a mammalian cell culture model, cleavage of the transmembrane protein TMEM67 by matrix metalloprotease ADAMTS9 regulates ciliogenesis (Ahmed et al., 2025). In *C. elegans*, the transmembrane proteoglycan syndecan plays a role in maintaining ciliary compartments (Acker et al., 2021) and matrix components secreted from glial cells protect neuronal cilia against damage (Varandas et al., 2025). In *Chlamydomonas*, ciliary polycystin protein PKD2 interacts with and organizes extracellular polymers called mastigonemes (Liu et al., 2020). It has also been proposed that cilia may respond directly to ECM via cilia-localized integrins (Pittman and Solecki, 2023). However, our understanding of how ECM impacts sensory cilia ultrastructure, protein localization, and function to influence transduction of sensory stimuli is incomplete.

*C. elegans* ciliated sensory neurons provide a model in which to study the relationship between cilia and ECM. The cilia of these neurons are housed within sense organs, which consist of one or more ciliated sensory neurons enwrapped by sheath and socket glial cells (Nechipurenko and Sengupta, 2025; Purice et al., 2024). Although most ciliated sense organs are found in the head of the worm, a few are found in the tail. The male has additional sex-specific sensory organs in the tail that mediate mate recognition and male mating behaviors (Barr and Garcia, 2006; Liu and Sternberg, 1995). Mating requires the cilia-localized TRP polycystin-2 channel protein PKD-2 (Barr et al., 2001), which is expressed by male-specific sensory neurons. PKD-2 also localizes to EVs, which are shed from the cilium in response to mechanical stimuli potentially from a mating partner (Wang et al., 2020). The sensory function and localization of PKD-2 to cilia and EVs is conserved from algae to humans (Hogan and Ward, 2024; Liu et al., 2023; Wang and Barr, 2018; Wood et al., 2013).

Here, we identify pleiotropic roles for the extracellular matrix gene *mec-9* in *C. elegans* ciliated sense organs. *mec-9* encodes a predicted secreted extracellular matrix protein containing Kunitz and EGF-like domains. *mec-9* was originally identified as a component of the ECM surrounding non-ciliated touch receptor neurons and whose mutation results in the *<u>mec</u>*hanosenory abnormal (Mec) phenotype (Du et al., 1996). In touch receptor neurons, MEC-9 organizes mechanosensory complexes composed of ion channels and specialized ECM structures (Das et al., 2024; Emtage et al., 2004). We find that neomorphic *mec-9(ok2853),* but not null *mec-9(my147)* or *mec-9(u437)* mutants, exhibit aberrant PKD-2::GFP accumulation at sensory cilia and defects in male mating behaviors. Consistent with a role for *mec-9* outside the touch receptor neurons, an isoform of *mec-9* of previously unknown function is expressed in ciliated sensory neurons in the head (inner and outer labial sense organs) and the male tail (hook and ray sense organs). In *mec-9(ok2853)* males, we observed accumulation of EVs in the cephalic sensory organ and defects in ciliary transition zone remodeling in inner labial type 2 (IL2) cilia. In the ray, labial, and cephalic sense organs, *mec-9* is not expressed in the EV producing neurons but rather is expressed by companion neurons. We propose a model in which *mec-9* acts cell non-autonomously from companion neurons to regulate sensory cilia function, ciliary protein localization, EV shedding, and ciliary ultrastructure.

## Results

### *mec-9* regulates PKD-2::GFP localization and male mating behavior

*C. elegans* male mating behaviors are mediated by male-specific sense organs found in the male tail. These include the ray sense organs, which consist of ciliated ray A-type (RnA) and B-type (RnB) neurons and a single glial cell, the ray structural cell (RnSt) (Figure 1A), and the hook sense organ, which consists of the ciliated HOA and HOB neurons and two glial socket and sheath cells (Sulston et al., 1980). The ray neurons appear to sense both chemical and mechanical stimuli (Barr et al., 2018; Koo et al., 2011; Luo et al., 2024; Zhang et al., 2018). Males also have sex-specific ciliated CEM neurons in the cephalic sensory organ. In males, this organ consists of the CEM and CEP neurons and two glial socket and sheath cells (Ward et al., 1975; Ware et al., 1975).

**Figure 1.**
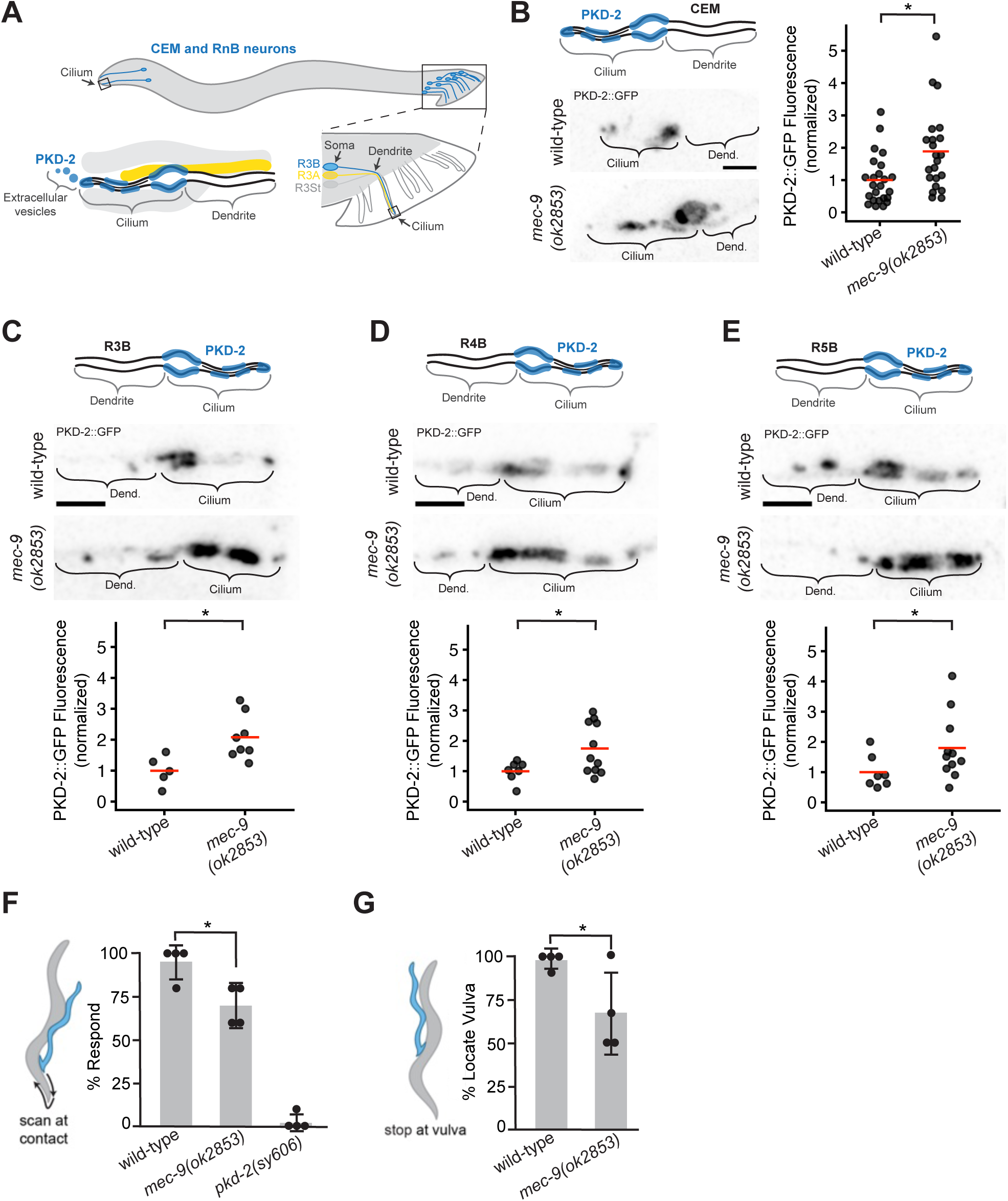
*mec-9(ok2853)* mutants exhibit PKD-2::GFP accumulation at cilia and defects in male mating behaviors. (A) Schematic of male-specific ciliated neurons and ray sensory organs in *C. elegans*. (B) Confocal images and quantification of PKD-2::GFP at CEM cilia in wild-type (PT3168) and *mec-9* mutants (PT4165). N ≥ 22 cilia. One-way ANOVA and post-hoc TukeyHSD, p = 0.01. In B-E, the cilia are oriented with anterior to the left and posterior to the right (in CEM cilia the tip is anterior, in RnB cilia the tip is posterior). (C, D, E) Confocal images and quantification of PKD-2::GFP at R3B cilia (C), R4B cilia (D) and R5B cilia (E) in wild-type and *mec-9* mutants. N ≥ 5 cilia. Two-way ANOVA revealed a significant main effect of genotype (p = 0.001) and no significant interaction between ray identity and genotype. For B-E, fluorescence values are normalized to the wild-type mean and red bars in dot plots indicate the mean. Scale bars = 2 µm. (F and G) *mec-9* mutants (PT4165) are defective in response to mating partner (F) and location of vulva (G) when compared to wild-type controls (PT3168). *pkd-2(sy606)* (PT9) is a negative control for mating assays. *pkd-2(sy606)* location of vulva is not shown because the number of responding males that can be scored for vulva location is very low. N = 4 mating assays, 10 animals per assay. One-way ANOVA and post-hoc testing with TukeyHSD, * = p < 0.01.

In wild-type animals, PKD-2 localizes to the ciliary base, shaft and tip of the male-specific CEM, RnB and HOB sensory neurons. In *mec-9(ok2853)* mutant males, PKD-2::GFP aberrantly accumulates at CEM cilia (Figure 1B). For the RnB cilia, we focused our analysis on R3B, R4B and R5B cilia because each cilium can be segmented for individual fluorescence quantification. R3B, R4B and R5B cilia all display aberrant PKD-2::GFP accumulation in *mec-9(ok2853)* mutants (Figure 1C-E).

Because proper PKD-2 localization is required for male mating behaviors, we asked if *mec-9(ok2853)* affects mating. When the ray neurons detect contact with a hermaphrodite, males respond by stopping forward locomotion and initiating backward scanning to search for the vulva (Barr and Garcia, 2006). *mec-9(ok2853)* males are defective in contact response, though not as severely as *pkd-2* mutants (Figure 1F). After responding, the male scans the hermaphrodite’s body to locate the vulva. Successful vulva location is defined as stopping upon encountering the vulva, and this behavior is mediated by the hook sense organ (Liu and Sternberg, 1995). *mec-9(ok2853)* males are also defective in vulva location (Figure 1G). Because of *mec-9*’s known role organizing mechanosensitive complexes in touch receptor neurons (Das et al., 2024; Du et al., 1996; Emtage et al., 2004), one possible interpretation of these results is that touch response contributes to male contact response and vulva location. However, *mec-4(e1611)* mutants, which are touch-insensitive due to loss of the touch receptor neurons (Chalfie and Sulston, 1981; Herman, 1987), do not exhibit defects in contact response or vulva location (Barr and Sternberg, 1999). We conclude that *mec-9* has other roles in addition to its function in touch receptor neurons, including in male-specific ciliated ray and hook sensory organs.

### The short isoform of *mec-9* is expressed in ciliated neurons and localizes to the distal dendrite region

*mec-9* encodes two isoforms: MEC-9L, the longer 839-amino acid isoform containing a signal peptide, 7 EGF-like domains, and 5 Kunitz domains, and MEC-9S, a shorter 502-amino acid isoform containing a signal peptide, 3 EGF-like domains, and 2 Kunitz domains (Figure S2A). Du, et al. (1996) showed that, in hermaphrodites, the long isoform is expressed by touch receptor neurons and the short isoform is expressed by a different group of unidentified neurons. *mec-9(ok2853)* is a 408-base pair in-frame deletion that removes 2 EGF-like domains, specifically the second and third EGF-like domains of the short isoform and the sixth and seventh EGF-like domains of the long isoform (Figure S2A). The defects in PKD-2::GFP localization and mating behaviors in *mec-9(ok2853)* mutants suggests that *mec-9* impacts the RnB neurons of the male tail (Koo et al., 2011). We thus wanted to investigate if either MEC-9 isoform is expressed in RnB neurons, in the companion RnA neurons, or in ray structural cells of the ray sensory organs. We examined long and short isoform expression in males using transcriptional reporters. Consistent with previous reports, we detected MEC-9L reporter expression in touch receptor neurons (Figure S1A). We detected MEC-9S reporter expression in cells of the male tail. MEC-9S reporter expression did not overlap with a marker for the RnB neurons (*P_klp-6_*::tdTomato) or the ray structural cells (*P_grl-18_*::mApple) (Cebul et al., 2020; Fung et al., 2023; Fung et al., 2020) (Figure 2A,B). Based on the morphology of these cells, specifically processes into the sensory rays which are most prominent in rays 2, 3, and 6, we conclude that MEC-9S is expressed in RnA neurons. We also detected MEC-9S reporter expression in the male-specific HOA neuron in the hook sense organ and in sex-shared IL1 and OLQ neurons of the inner and outer labial sense organs (Figure 2C,D). Our results are consistent with available single-cell and single-nucleus RNA-Seq data (Morillo et al., 2025; Olson; St Ange et al., 2025; Taylor et al., 2021; Weinreb et al., 2025) (Figure S1C). Because MEC-9S is expressed by RnA but not RnB neurons, we conclude that *mec-9* acts cell non-autonomously to regulate PKD-2 ciliary localization in ray sensory organs.

**Figure 2.**
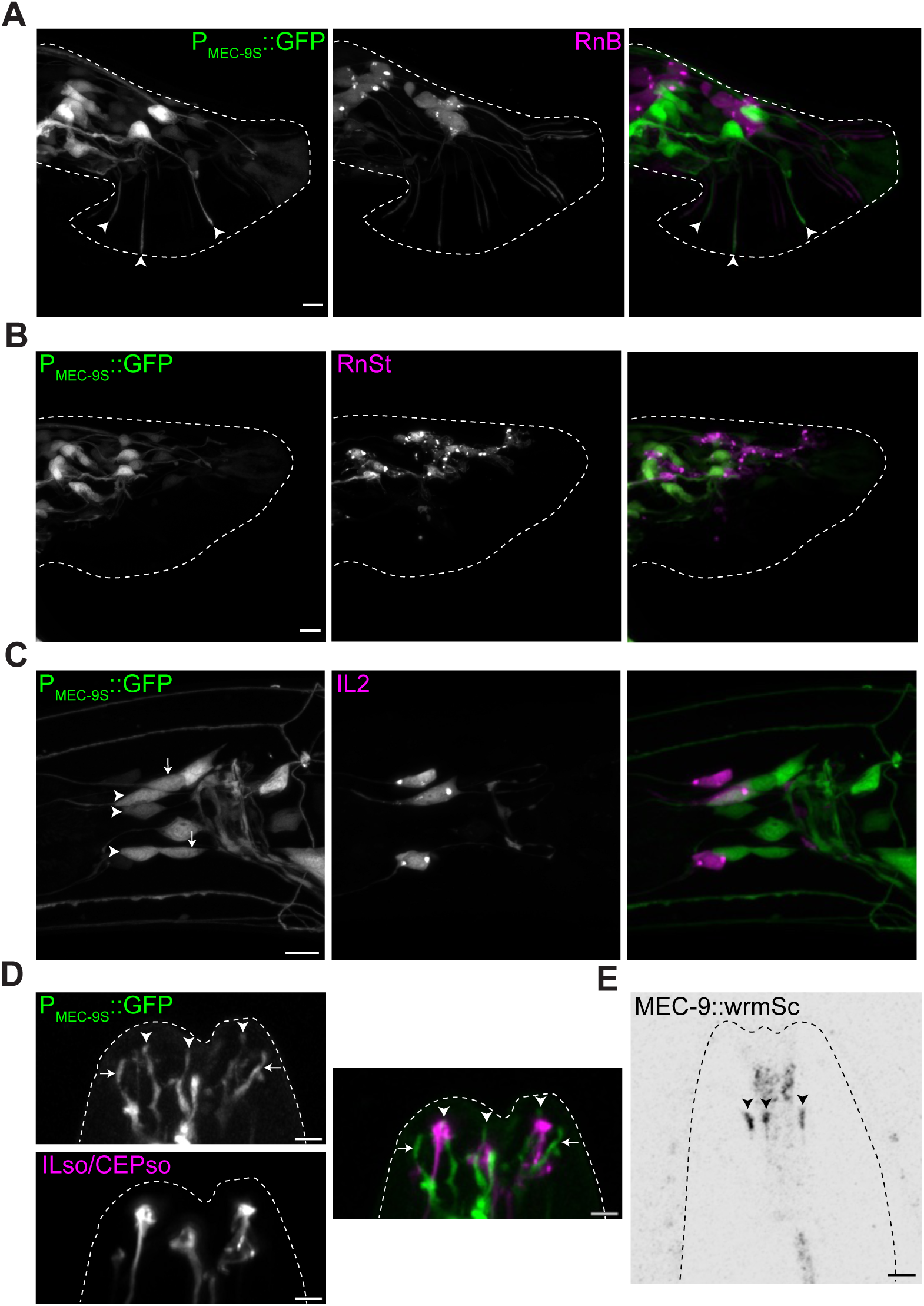
The short isoform encoded by *mec-9* is expressed in the male-specific RnA neurons and in the sex-shared IL1 and OLQ neurons. (A) Confocal image of MEC-9S transcriptional reporter (green) in adult male tail, co-expressed *with P_klp-6_::tdTomato* (magenta) which labels RnB neurons (PT4357). Arrowheads: RnA neurons. Scale bar = 5 µm. (B) Confocal image of MEC-9S transcriptional reporter (green) in adult male tail co-expressed with *P_grl-18_::mApple* (magenta) which labels RnSt cells (Fung et al., 2023; Fung et al., 2020). Scale bar = 5µm. The *P_grl-18_::mApple* reporter causes a Male ABnormal (Mab) phenotype in which the sensory rays do not form properly, this is why no MEC-9S reporter signal is detected in rays. (C) Confocal image of MEC-9S transcriptional reporter (green) around nerve ring in L4 hermaphrodite, co-expressed with *P_klp-6_::tdTomato* (magenta) which labels IL2 neurons (PT4357). Arrows: OLQ. White arrowheads: IL1. Scale bar = 5 µm. (D) Confocal image of MEC-9S transcriptional reporter (green) in head sensilla in adult male, co-expressed with *P_grl-18_::mApple* (magenta) which labels ILso and CEPso cells (PT4356) (Cebul et al., 2020; Fung et al., 2023). Arrows: OLQ. White arrowheads: IL1. Scale bar = 2 µm. (E) Confocal image of endogenously-tagged MEC-9::wrmScarlet in L4 male head sensilla (PT4128). Arrowheads: IL1 distal dendrite region. Scale bar = 2 µm.

To determine the subcellular localization of MEC-9S, we examined an endogenous translational fusion reporter designed to tag both isoforms of *mec-9* at the C-terminus with wrmScarlet (Das et al., 2024). In the male tail, we only detected long isoform expression in the PLM neurons with this reporter (Figure S1B). In adult hermaphrodites and males, we detected fluorescence in the distal dendrite region of IL1 neurons (Figure 2E). Our transcriptional reporter indicated that MEC-9S is expressed in the IL1 neurons, suggesting that this fluorescence represents the subcellular localization of MEC-9S. The MEC-9S transcriptional reporter revealed a broader expression pattern than the endogenous fusion reporter, which could be due to difficulties in detecting secreted fluorescent proteins in the extracellular space and/or low levels of expression (Heppert et al., 2016).

### *mec-9(ok2853)* is a neomorphic allele

A mutation impacting only MEC-9L, *mec-9(e1494)* (Figure S2A), does not affect contact response or vulva location behaviors (Barr and Sternberg, 1999). Furthermore, the *mec-4(e1611)* mutation, which results in loss of the touch receptor neurons and therefore loss of MEC-9L secreted by these neurons, does not affect contact response or vulva location (Barr and Sternberg, 1999). These previous results, combined with the expression patterns of MEC-9L and MEC-9S, led us to hypothesize that the defects in PKD-2::GFP localization and male mating behavior in *mec-9(ok2853)* mutants are due to a short-isoform-specific function.

To test this hypothesis, we generated a STOP-IN allele, *mec-9(my147)*, specifically disrupting the short isoform at its unique N-terminus (Figure S2A). *mec-9(my147)* introduces an early stop codon in the exons encoding MEC-9S and does not alter those encoding MEC-9L. Interestingly, *mec-9(my147)* mutants display the PKD-2::GFP ciliary accumulation specifically in R5B (Figure S2B-E), but not other ray neurons or CEMs. However, *mec-9(my147)* mutants do not exhibit defects in male mating behaviors (Figure S2F-G). There are multiple ways to interpret these results. One possibility is that MEC-9L and not MEC-9S has a role in ray sensory neurons. However, we previously showed that a MEC-9L-specific mutant does not exhibit mating defects, arguing against a role for the long isoform (Barr and Sternberg, 1999). A second possibility is that MEC-9L and MEC-9S play redundant roles and the *mec-9(ok2853)* allele impacts the function of both isoforms. We tested an additional null allele, *mec-9(u437)*, which has an 11-base pair frame-shift deletion predicted to affect both isoforms. *mec-9(u437)* mutants do not exhibit defects in male mating behaviors (Figure S3A-B). The third possibility is that *mec-9(ok2853)* is a neomorphic allele that causes a phenotype not seen in nulls. Most neomorphic alleles are dominant, so we determined if *mec-9(ok2853)* is dominant or recessive by assaying male mating behavior in heterozygotes. Heterozygous males do not exhibit defects in male mating behavior (Figure S3C-D). Thus, *mec-9(ok2853)* appears to be a recessive, neomorphic allele. To test the possibility that mating defects seen in *mec-9(ok2853)* animals are due to a mutation elsewhere in the genome, we generated the identical deletion via CRISPR-Cas9 genome editing in wild-type animals. This new allele, *mec-9(syb11045)*, resulted in similar defects in contact response and vulva location (Figure S3E-F).

### External environmental EV release is not impaired in *mec-9(ok2853)* mutants

Cilia shed EVs from two locations, the ciliary tip and the ciliary base (Wang et al., 2021). Ciliary tip EVs are released into the external environment, while ciliary base EVs are released into the ECM-filled lumen formed by the glial cells. Previously identified mutants that display PKD-2::GFP ciliary accumulation have shown either a reduction or no change in the amount of PKD-2::GFP-labeled EVs shed into the environment from the ciliary tip (Akella et al., 2020; Maguire et al., 2015; O’Hagan et al., 2017; Silva et al., 2017; Wang et al., 2014). We therefore asked if *mec-9(ok2853)* mutants have a defect in ciliary tip shedding of EVs into the environment by measuring environmental PKD-2::GFP EV release from *mec-9(ok2853)* male rays.

EVs are dynamically released in response to confinement under a coverslip, such that the time the animal spends on the imaging slide affects the number of EVs shed into the environment (Wang et al., 2020; Wang et al., 2024b). To account for this, we imaged PKD-2::GFP-labeled EVs immediately after mounting (Figure 3A, 0 h) and then imaged the same animal after 1 hour (Figure 3A, 1 h). Wild-type animals display an increase in the number of PKD-2-positive EVs released from 0 h to 1 h (Figure 3B). Similar to wild-type animals, *mec-9(ok2853)* displayed increases in PKD-2-positive EV numbers from 0 h to 1 h (Figure 3B). We conclude that environmental ciliary EV release is not impaired in *mec-9(ok2853)* mutants.

**Figure 3.**
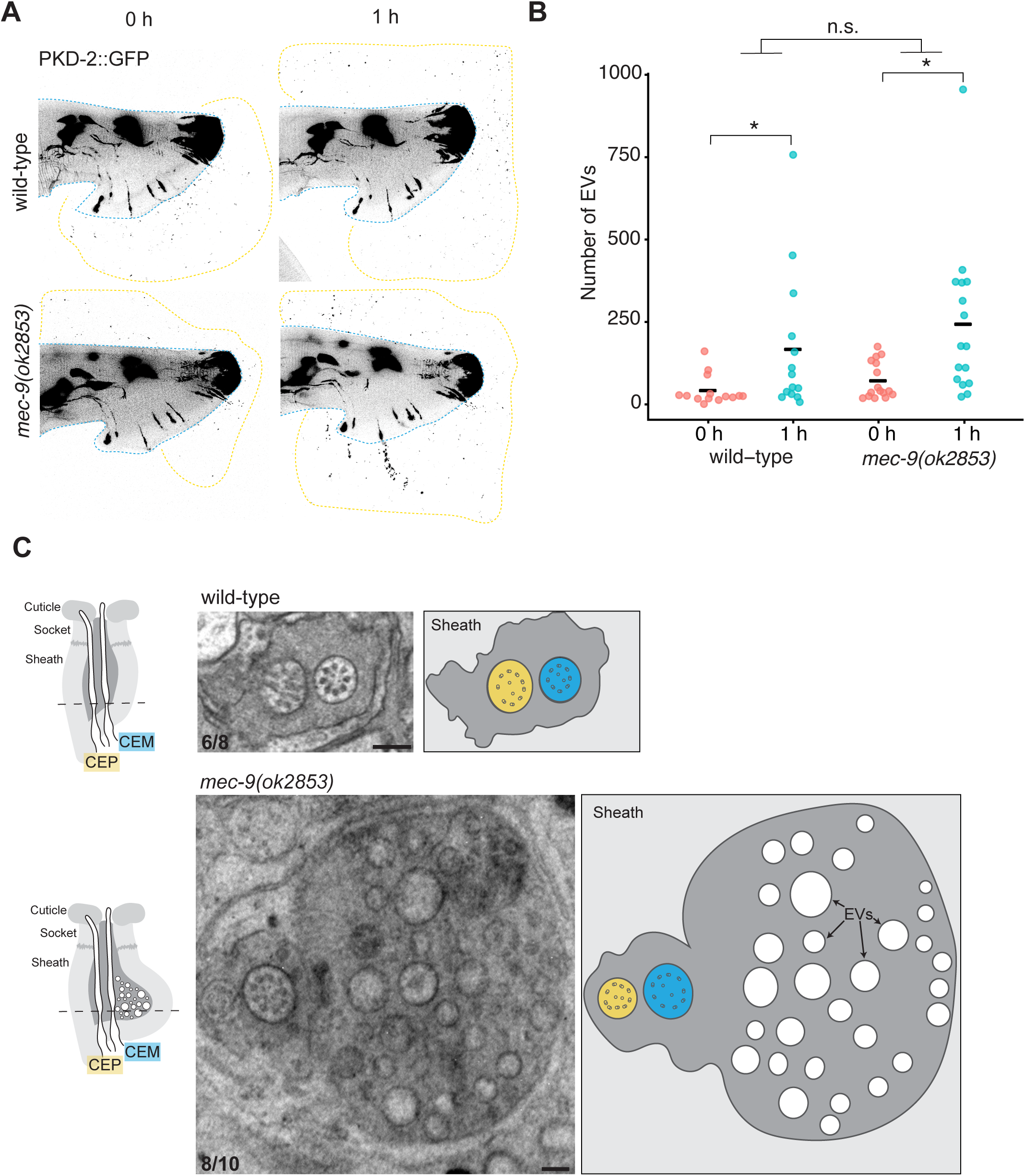
*mec-9(ok2853)* mutants exhibit no defect in externally-released PKD-2::GFP EVs but display increased EVs in the lumen of the cephalic sensory organ. (A) Representative confocal images of wild-type (PT3168) and *mec-9(ok2853)* (PT4165) male tails just after mounting on slide (0 h) and after 1 hour (1 h). The blue dashed lines outline the tail, the yellow dashed lines enclose regions containing externally-released PKD-2::GFP EVs. (B) Quantification of PKD-2::GFP EVs. Each data point represents the total EV count released by an individual male, black bars indicate the mean (N = 14 males per genotype). Repeated measures ANOVA revealed no significant (p<0.05) effect of genotype. (C) TEM cross sections of the cephalic sensory organ at the level of the ciliary transition zone in wild-type and *mec-9(ok2853)* mutants. N = 8-10 sense organs examined across 2-3 animals per genotype. Numbers in lower left corner of images indicate the number of sense organs exhibiting the phenotype shown over the total number of sense organs examined. Scale bars = 200 nm. In the cartoons, neurons are the colors indicated, the sheath cell is light gray, and the lumen is dark gray.

### *mec-9(ok2853)* causes internal EV accumulation and impacts cephalic sensory organ morphology

To ask if *mec-9(ok2853)* affects EV release from the ciliary base, we used transmission electron microscopy to examine cephalic sensory organs in young adult *mec-9(ok2853)* mutants. In males, the cephalic sense organ contains the cilia of the CEP and CEM neurons enwrapped by sheath and socket glia, with CEM cilia protruding through a cuticular pore into the environment. Strikingly, the lumen formed by the glial cells was greatly expanded and full of EVs in *mec-9(ok2853)* mutants (Figure 3C). This phenotype is reminiscent of other mutants that display PKD-2::GFP accumulation at cilia, increased EVs within an expanded lumen, and external environmental EV shedding defects (Akella et al., 2020; Maguire et al., 2015; O’Hagan et al., 2017; Silva et al., 2017; Wang et al., 2014). *mec-9(ok2853)* mutants do not affect external environmental EV shedding of PKD-2::GFP (Figure 3A, B) and do not display ultrastructural defects in CEM cilia (Figure 3C).

PKD-2 is a cargo of ciliary EVs shed from the CEM ciliary tip and base, so one possible explanation for the increased PKD-2::GFP signal at the ciliary base is accumulation of cilia base EVs in the lumen. Our transcriptional reporter for MEC-9S did not show expression in the CEM or CEP neurons of the cephalic sensory organ, but did show expression in the adjacent OLQ neuron. The distal dendrites of the OLQ and CEM neurons are in close proximity (Ward et al., 1975) and OLQ forms synaptic connections with both CEM and CEP (Cook et al., 2019; Emmons, 2024). Similar to our finding that *mec-9* acts cell non-autonomously in the ray sensory organs, these data suggest that the cephalic sensory organ phenotypes may be caused by cell non-autonomous action of *mec-9*.

### *mec-9(ok2853)* impacts inner labial sensory organs and IL2 ciliary microtubule remodeling

Based on the expression pattern of the MEC-9S reporter and localization of endogenous MEC-9::wrmScarlet in IL1 neurons (Figure 2C-E), we examined the ultrastructural anatomy and morphology of inner labial (IL) sense organs in young adult males. These sense organs contain the cilia of two neurons, IL1 and IL2, enwrapped by two glial cells, the IL sheath and socket (Doroquez et al., 2014). The base of the IL1 cilium has a canonical transition zone (TZ) with 9 doublet microtubules (dMTs). The TZ of IL2 cilia undergoes developmental remodeling to shift from containing the canonical 9 dMTs in early larval stages to ∼ 6 dMTs in adulthood (Akella et al., 2019; Doroquez et al., 2014; Ward et al., 1975). We did not observe any defects in glial cell morphology or IL1 ciliary ultrastructure in the IL sense organs of *mec-9(ok2853)* mutants (Figure 4A). However, unexpectedly, the IL2 TZs of *mec-9(ok2853)* mutants do not fully complete the remodeling process and display 6-9 dMTs compared to 5-7 dMTs in wild-type (Figure 4A-B). These results suggest that *mec-9(ok2853)* impacts processes involved in IL2 ciliary transition zone remodeling and are consistent with *mec-9* acting cell non-autonomously.

**Figure 4.**
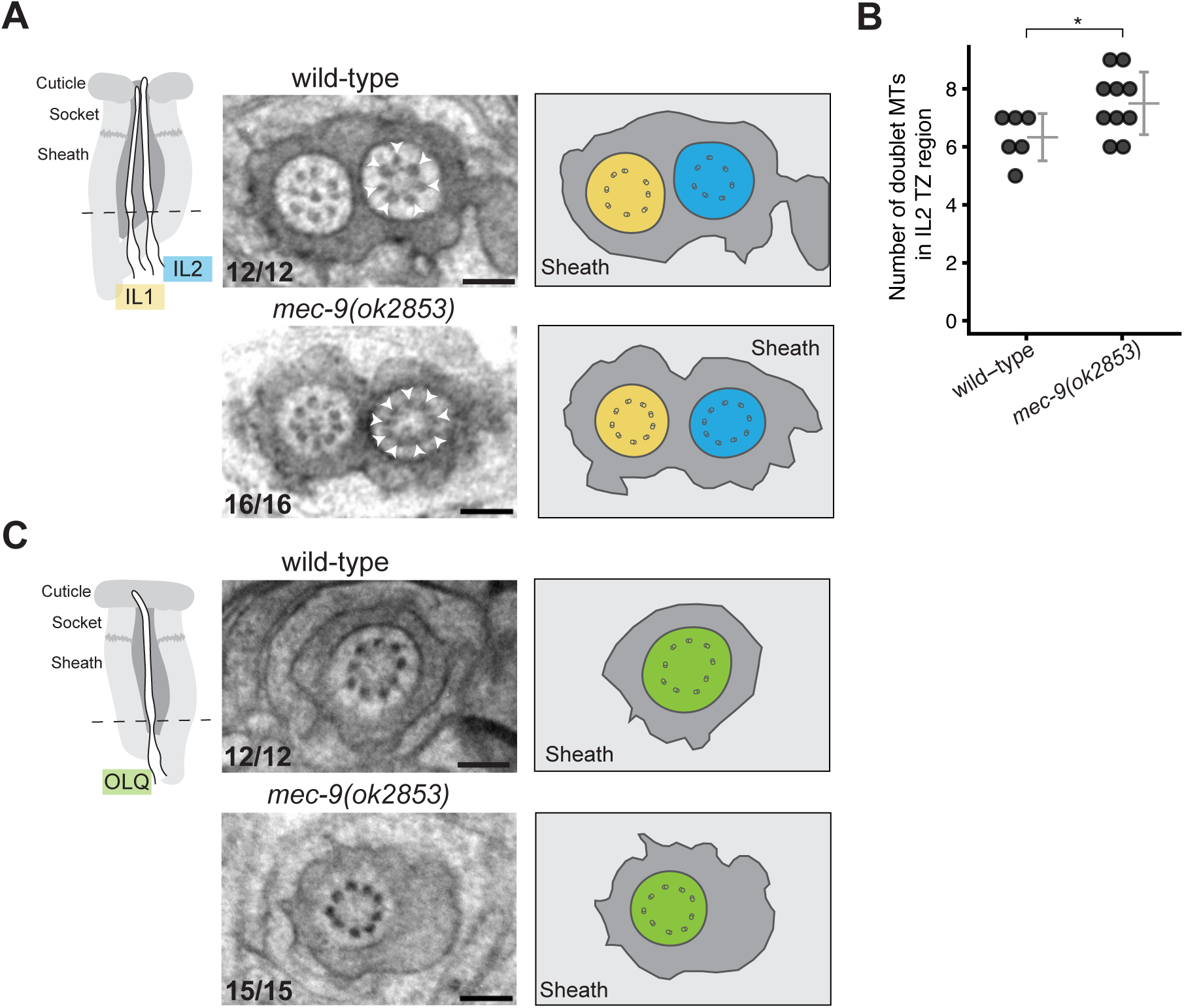
*mec-9(ok2853)* impacts IL2 ciliary transition zone remodeling. Dashed lines in cartoons denote level of TEM cross sections (ciliary transition zone). (A-B) The IL2 cilia of *mec-9(ok2853)* mutants exhibit an increased number of doublet microtubules (white arrowheads) in the transition zone (TZ) compared to wild-type. IL1 cilia ultrastructure and the lumen of the IL sensory organ appear morphologically normal. N = 12-16 sense organs examined across 2-3 animals per genotype. For quantification in (B), N ≥ 6 TZs per genotype. T-test p = 0.04. Gray bars in dot plot indicate the mean +/- SD. (C) Outer labial sensory organs appear morphologically normal in *mec-9(ok2853)* mutants. N = 12-15 sense organs examined across 2-3 animals per genotype. Numbers in lower left corner of images indicate the number of sense organs exhibiting the phenotype shown over the total number of sense organs examined. Scale bars = 200 nm. In the cartoons, neurons are the colors indicated, the sheath cell is light gray, and the lumen is dark gray.

To further investigate the effect of *mec-9(ok2853)* on IL2 cilia, we examined a fluorescent reporter for CIL-7. CIL-7 is a myristoylated protein that localizes to IL2, CEM and RnB cilia and is required for PKD-2 localization to cilia and for environmental shedding of PKD-2-laden EVs (Maguire et al., 2015; Wang et al., 2021). We observed aberrant accumulation of CIL-7::GFP in *mec-9(ok2853)* IL2 cilia (Figure S4). We also observed aberrant accumulation of CIL-7::GFP in *mec-9(ok2853)* CEM cilia (Figure S4), reminiscent of the PKD-2::GFP accumulation we observed in CEM cilia. We conclude that *mec-9(ok2853)* causes defects in IL2 cilia TZ remodeling and CIL-7 ciliary localization. Because *mec-9S* is expressed by IL1 but not IL2 neurons and MEC-9::wrmScarlet localizes to IL1 distal dendrites, this is consistent with cell non-autonomous action of *mec-9*.

Based on our observation of MEC-9S reporter expression in OLQ neurons (Figure 2D), we examined the ultrastructural anatomy and morphology of the outer labial quadrant sense organs in *mec-9(ok2853)* mutants. The mechanosensory OLQ sense organs consist of one ciliated neuron and two glial cells. We did not observe any ultrastructural or morphological defects in the ciliary transition zone or glial cells of OLQ sense organs in *mec-9(ok2853)* mutants (Figure 4C).

### *mec-9(ok2853)* impacts amphid sensory organ morphology

In the course of our TEM analysis, we noted morphological defects in the amphid sensory organs of *mec-9(ok2853)* mutants. In wild-type animals, 10 amphid channel cilia are enclosed by the lumen formed by amphid sheath and socket glia (Figure S5A). The glial cells are thought to secrete extracellular matrix material that surrounds the cilia (Doroquez et al., 2014; Perkins et al., 1986). In *mec-9(ok2853)* mutants, fewer cilia populate the lumen. Specifically, the cilia of the amphid neurons that possess two cilia each (ADF and ADL) do not extend completely in the lumen, while amphid channel neurons with a single cilium appear normal (Figure S5B). The matrix-filled lumen formed by the amphid glial cells is expanded in *mec-9(ok2853)* mutants (Figure S5B).

To further investigate the effect of *mec-9(ok2853)* on amphid cilia, we examined an endogenous fluorescent reporter for tetraspanin TSP-6, a transmembrane protein that localizes to amphid and phasmid cilia and ciliary EVs (Nikonorova et al., 2022; Razzauti and Laurent, 2021). The TSP-6::mNG localization pattern was similar in *mec-9(ok2853)* mutants and wild-type controls, with no detectable differences in amphid ciliary membrane morphology in the head or phasmid cilia length in the tail (Figure S5C-E). We conclude that *mec-9(ok2853)* impacts amphid sensory organ morphology, but not TSP-6 localization to cilia. Because we did not detect MEC-9S reporter expression in amphid neurons or glia, the effect on amphid sensory organ morphology appears to be cell non-autonomous.

## Discussion

The functioning nervous system depends on interactions between neurons and their companion cells. Here, we show that *mec-9* is expressed in companion neurons and acts non-autonomously to regulate ciliary protein localization and EV shedding in cephalic, labial, and ray sense organs in the adult *C. elegans* male. Using a combination of behavioral analysis, live super-resolution imaging, and transmission electron microscopy, we found that *mec-9(ok2853)* male mutants display mating defects (Fig. 1F-G), mislocalization of ciliary proteins PKD-2 and CIL-7 in CEM, RnB, and IL2 neurons (Fig. 1B-E, S4), accumulations of EVs within cephalic organs (Fig. 3C), and abnormal ultrastructure of the IL2 ciliary transition zone (Fig. 4A-B). *mec-9* expression is not detectable in these neurons, but is evident in companion neurons: the OLQ neurons (closely associated with cephalic organs), RnA neurons (associated with RnBs in the ray organs), and IL1 neurons (associated with IL2s in the inner labial organs) (Fig. 2, S1C). Combined, these data indicate that companion neurons act non-autonomously to influence the properties and functions of nearby ciliated neurons (Figure 5).

**Figure 5.**
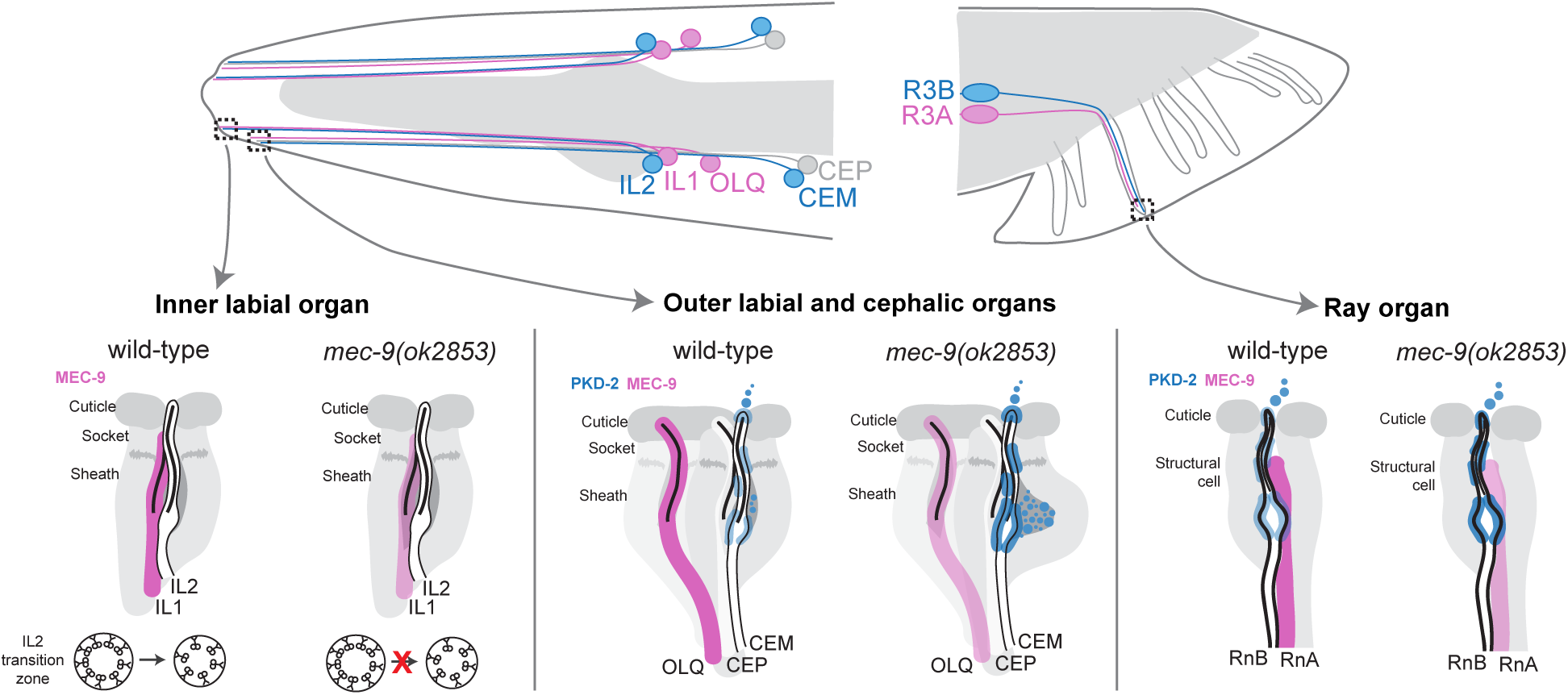
Model: *mec-9* acts cell non-autonomously from companion neurons (magenta) to regulate cilia structure and function in CEM, IL2 and RnB neurons (blue). In the outer labial and cephalic sensory organs, the short isoform encoded by *mec-9* is expressed by OLQ neurons and *mec-9(ok2853)* causes defects in PKD-2 localization and EV accumulation at the ciliary base of CEM neurons. In the inner labial sensory organs, the short isoform encoded by *mec-9* is expressed by IL1 neurons and *mec-9(ok2853)* causes defects in IL2 ciliary TZ remodeling. In the ray sensory organs, the short isoform encoded by *mec-9* is expressed by RnA neurons and *mec-9(ok2853)* causes defects in male mating behavior and PKD-2 accumulation at RnB cilia.

### Neuron-to-neuron cell non-autonomous activity of *mec-9*

We detected PKD-2::GFP accumulation in RnB cilia in *mec-9(ok2853)* mutants (Figure 1C-E), however we did not detect *mec-9* expression in these cells. This may reflect a technical limitation of our reporters (Figure 2); however, our reporter results agree with available single-cell RNA sequencing data in which *mec-9* expression is highest in RnA neurons (Morillo et al., 2025; Olson) (Figure S1C). Therefore, we propose that *mec-9* acts cell-non-autonomously. What might explain cell non-autonomous activity of *mec-9*? In the ray sensory organs, one possible cell non-autonomous mechanism of action is through the shared ECM surrounding the neurons: changes in secreted ECM components from RnA neurons could impact adjacent RnB neurons to cause accumulation of PKD-2.

In the cephalic organ, PKD-2::GFP accumulated at the CEM ciliary base (Figure 1B) and EVs accumulated in the cephalic lumen of *mec-9(ok2853)* males (Figure 3C). However, we did not detect *mec-9* expression in the companion CEP neurons in the cephalic sensory organ. In this case, *mec-9* appears to act cell non-autonomously between sensory organs rather than within a sensory organ. How might this occur? The distal dendrites of the *mec-9*-expressing OLQ neurons and the CEM neurons are in close proximity (Ward et al., 1975). Thus, though these neurons occupy separate sensory organ compartments more anteriorly, OLQ and CEM neurons share ECM at the level of the distal dendrite and we propose that *mec-9* may act cell non-autonomously through this shared ECM.

In the inner labial sense organ, *mec-9(ok2853)* mutants exhibit defects in the developmental remodeling of the transition zone of IL2 cilia (Figure 4A-B). In wild-type animals, the IL2 TZ developmentally remodels from containing the canonical 9 doublet microtubules in young larvae to non-canonical 6 doublet microtubules in adulthood (Akella et al., 2019; Doroquez et al., 2014; Perkins et al., 1986). *mec-9(ok2853)* IL2 TZs contain 7-8 doublet microtubules in adulthood, suggesting they do not complete remodeling. *mec-9* is expressed by the companion IL1 neuron in the inner labial organ, suggesting cell non-autonomous action via shared ECM. Other known factors that influence IL2 transition zone remodeling are *che-13* (IFT57), *osm-1* (IFT72), *osm-5* (IFT88), *osm-6* (IFT52), *rab-28* (ciliary GTPase) and *tba-6* (alpha-tubulin) (Akella et al., 2023; Perkins et al., 1986). With the exception of *mec-9*, all are thought to act cell autonomously in IL2 cilia. Thus, our findings suggest that companion cells can influence ciliary ultrastructure of nearby cells. Mammalian cilia exhibit diverse microtubule organization (Ott et al., 2024), thus an understanding of non-autonomous factors governing ciliary specialization is likely to be relevant to human diseases such as ciliopathies.

Cell non-autonomous mechanisms, including ECM, play important roles in the nervous system (Coraggio et al., 2024; DeVault et al., 2018; Filipczak et al., 2025; Hansen and Hippenmeyer, 2020). Glia are thought to be major producers of ECM in the nervous system in mammals and *C. elegans* (Fung et al., 2024), and glial secretion is altered in neurodevelopmental disorders (Caldwell et al., 2022). Neurons also contribute to ECM, and neuronal ECM can have cell non-autonomous effects. For example, lumican, an extracellular proteoglycan, is secreted by one subpopulation of corticospinal neurons to prevent innervation by a different subpopulation of corticospinal neurons (Itoh et al., 2023). Our work suggests that altering ECM secretion by neurons (in this case, of abnormal MEC-9) can have non-autonomous, deleterious effects throughout the nervous system.

### Neomorphic function of *mec-9(ok2853)*

We show that *mec-9(ok2853)* acts as a recessive neomorphic allele, causing phenotypes that are not observed with null alleles. Rather than eliminating gene function, as in a null, *mec-9(ok2853)* retains an intact reading frame that could produce a novel protein product, which could act in different ways to cause the observed pleiotropic phenotypes. So why study neomorphic alleles that are difficult to interpret? Without studying a neomorphic allele, we would not have uncovered a role for *mec-9* in ciliated sensory organs. Only ∼10% of the 180 cuticular collagens of *C. elegans* were identified by phenotypic screening, which may reflect genetic redundancy of ECM genes (Li Zheng et al., 2020). Furthermore, in humans, null mutations in ECM genes result in less severe disease than missense or in-frame exon-skipping mutations (Lamandé and Bateman, 2020). This difference in disease severity is thought to occur because null mutations may be compensated for with functional redundancy, while structural mutations can result in misfolded protein that triggers ER stress inside the expressing cells in addition to altering the ECM. We do not know if or how *mec-9(ok2853)* impacts other ECM components of ciliated sensory organs.

### Potential mechanosensitive role of *mec-9* in IL1, OLQ, and RnA neurons

In touch receptor neurons, MEC-9, along with ECM components MEC-1 and MEC-5, is required for the formation of mechanosensory complexes containing nidogen and laminin (Das et al., 2024). An interesting possibility is that MEC-9S plays a mechanosensory role in ciliated sensory neurons. The IL1 and OLQ neurons sense nose touch and regulate foraging movements (Hart et al., 1995; Kaplan and Horvitz, 1993). Interestingly, the ciliated neurons that express *mec-9S* (including IL1, OLQ, and RnA) have long striated rootlets, structures that extend from the base of the cilium into the dendrite and may provide support for resistance to mechanical stress or support mechanosensation (Doroquez et al., 2014; Sulston et al., 1980). It will be interesting to identify the ECM components of ciliated sensory organs and to investigate their role in mechanosensation in future studies.

### ECM and ciliopathies

ECM components affect sensory cilia, critical organelles whose function underlies a class of human diseases termed ciliopathies. Chick primary cilia contain ECM receptors such as integrins (McGlashan et al., 2006), and matrix metalloproteases can cleave transmembrane ciliary proteins (Ahmed et al., 2025). In vertebrates, the ECM protein Eyes shut is necessary for ciliary pocket morphology and photoreceptor survival (Yu et al., 2016), with mutation causing photoreceptor degeneration in retinitis pigmentosa patients (Abd El-Aziz et al., 2008). In *C. elegans*, the ECM component DYF-7 regulates amphid, CEP, OLQ and IL2 dendrite extension, which then affects cilia placement (Heiman and Shaham, 2009; Low et al., 2019), and mutation of *mec-1*, which encodes an ECM component, results in ciliary fasciculation defects of amphid sensory neurons (Perkins et al., 1986). We extend these findings about the role of ECM at cilia to include regulating polycystin localization, microtubule ultrastructure, and EV shedding from the ciliary base. Because abnormal ECM and fibrosis are hallmarks of ciliopathies, we propose that feedback between cilia and ECM may contribute to pathogenesis in ciliopathies.

### Limitations of the study

We were unable to determine the subcellular localization pattern of MEC-9 protein in the outer labial, cephalic, and ray sensory organs using the endogenous *mec-9::wrmScarlet* translational fusion reporter and various extrachromosomal transgenes with *mec-9* fused to fluorescent protein (De Vore, 2018). Alternative fluorophores or changing the location of the tag could improve visualization.

## Methods

### *C. elegans* culture and maintenance

Nematodes were maintained using standard conditions (Brenner, 1974). Males and hermaphrodites were isolated at the 4^th^ larval L4 stage 24hrs prior to experiments and kept at 20°C overnight. In *C. elegans*, the predominant sex is hermaphrodite and males spontaneously arise only rarely (less than 1%). Therefore, in all experiments in which males were tested, we used strains with either the *him-5(e1490)* or *him-8(e1489)* allele as wild type. *C. elegans* sex is determined by X to autosome ratio, with hermaphrodites being XX and males being hemizygous for X. High incidence of male (Him) gene 5 (*him-5*) mutations increase the rate of X chromosome non-disjunction (Hodgkin et al., 1979). *him-5(e1490)* and *him-8(e1489)* males exhibit normal mating behaviors and are used as positive controls for mating assays (Walsh et al., 2020).

### Generation of strains

Strains used in this study are listed in Table 1. *mec-9(ok2853)* (Consortium, 2012), *myIs1* PKD-2::GFP (Bae et al., 2006), *mec-9(u437)* (Du et al., 1996) (gift from Chalfie Lab, Columbia University), *pkd-2(sy606)* (Barr et al., 2001), *hmnIs47* grl-18p::mApple (Fung et al., 2020) (gift from Heiman Lab, Harvard Medical School/Boston Children’s Hospital), *pg143*[mec-9::wSc::Lox2272::3xMyc] (Das et al., 2024), (*him-5(e1490)* (Hodgkin, 1983), and *him-8(e1489)* (Hodgkin, 1983) described previously. Strains arising from mutagenesis screens were backcrossed at least 6 times.

**Table 1.**
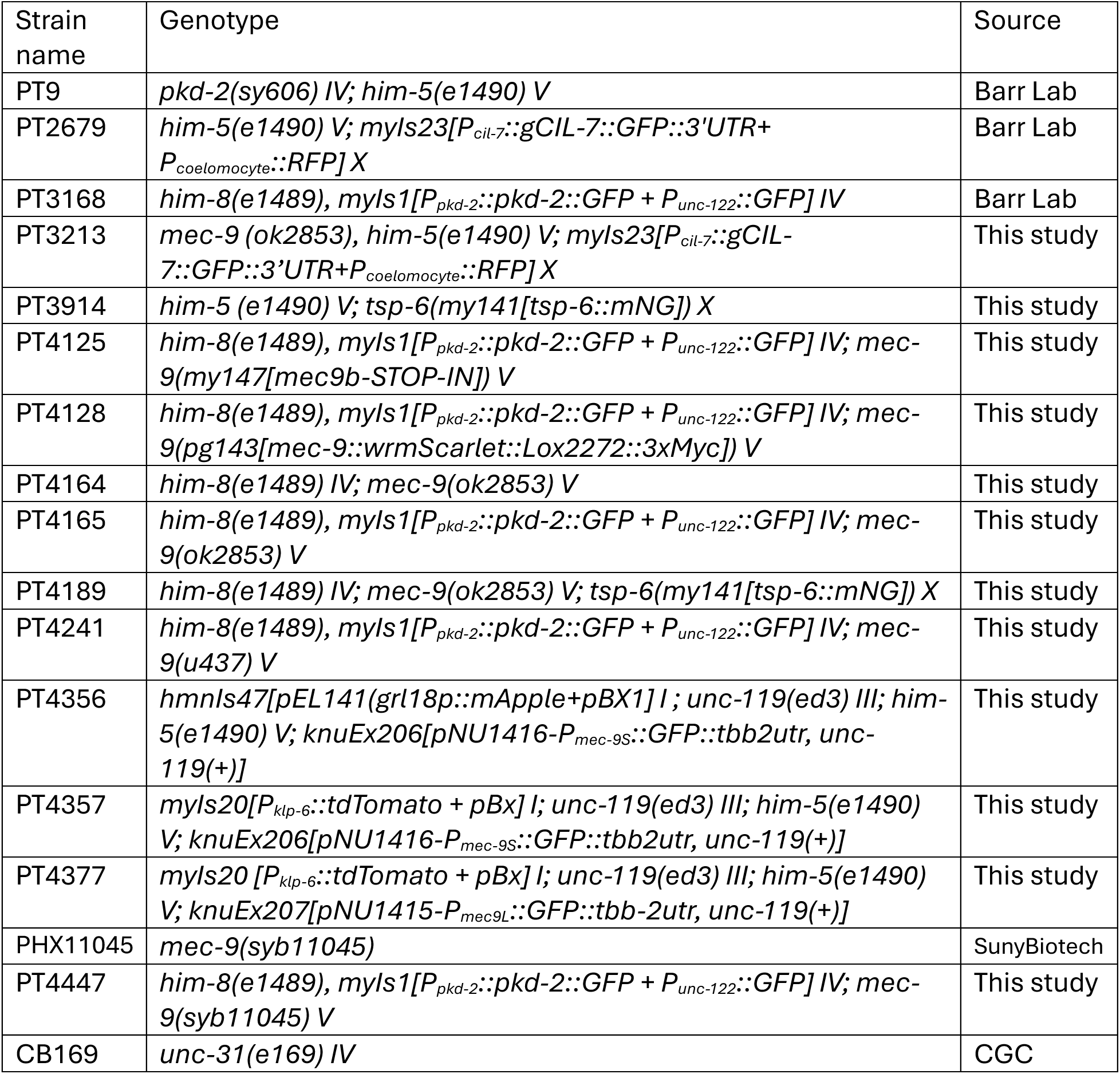
Strains used in this study.

The transcriptional reporters for *mec-9S* and *mec-9L* were constructed by Knudra Transgenics. To create the *mec-9L* reporter, Knudra generated a plasmid containing the *mec-9L* promoter region (the 2 kb upstream of the *mec-9L* reading frame) driving GFP followed by the *tbb-2* 3’ UTR. For the *mec-9S* reporter, Knudra generated a plasmid containing the *mec-9S* promoter region (1.5 kb upstream of the *mec-9S* reading frame, this region is within the 9^th^ intron of the full *mec-9* gene) driving GFP and followed by the *tbb-2* 3’ UTR. Both plasmids contain an *unc-119(+)* selection cassette. *unc-119* worms were injected and coordinated animals that carry the reporter were selected for maintenance of the constructs as extrachromosomal arrays.

To create the *mec-9(my147)* mutant and *tsp-6(my141[tsp-6::mNG])* via CRISPR, we followed the protocol of Dokshin et al. (Dokshin et al., 2018). Guide sequences were designed with CRISPOR (Haeussler et al., 2016), and we introduced silent mutations into the PAM site (Table 2). *mec-9(my147)* introduces early STOP codons into the first exon of *mec-9S* (Wang et al., 2018). The STOP-IN cassette also includes an Nhe1 site to facilitate genotyping. CRISPR-generated strains were validated with Sanger sequencing and backcrossed at least 3 times.

**Table 2.**
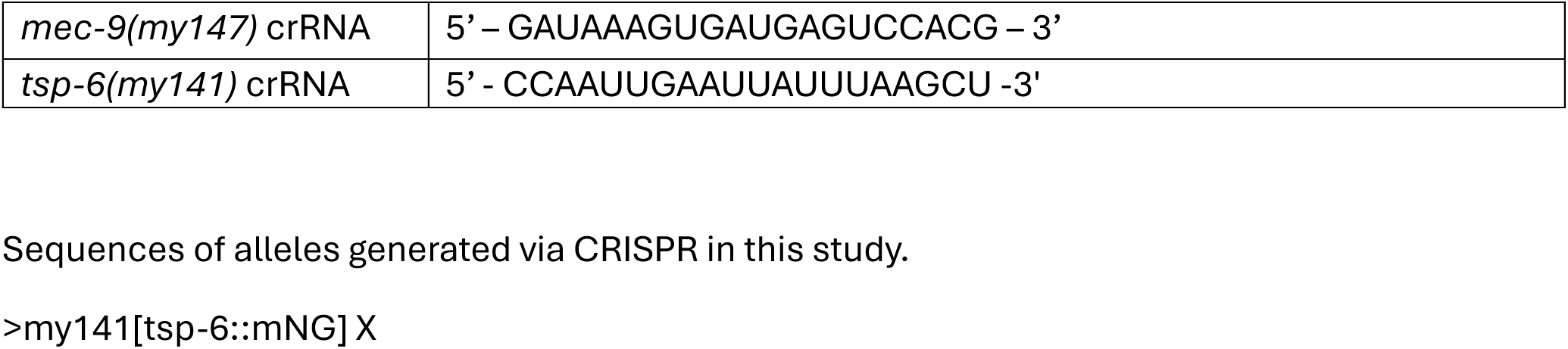
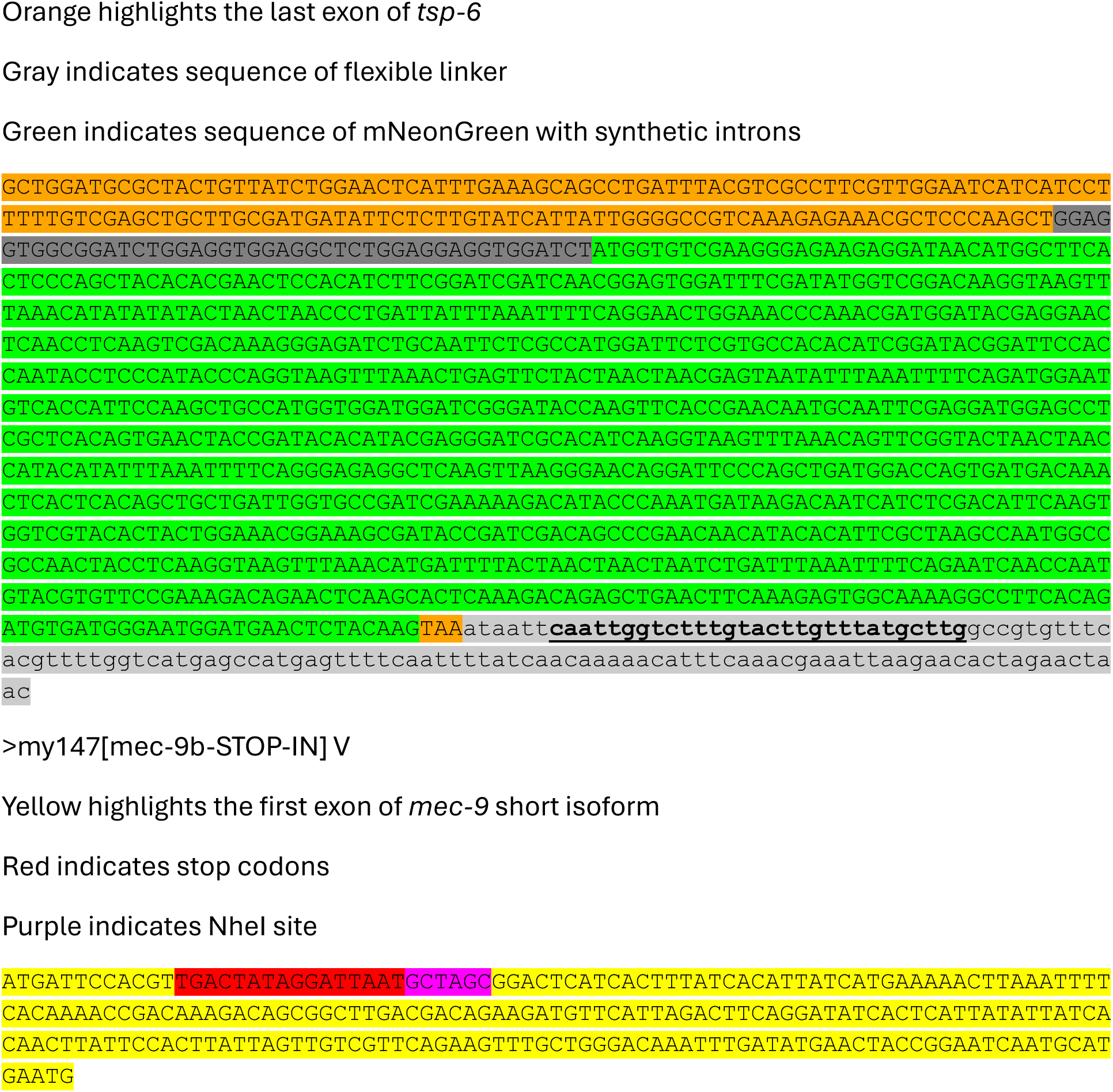
crRNAs for Cas-9-mediated genome editing.

*mec-9(syb11045)* was created by SunyBiotech in N2 background and deletes identical 408 bp as *mec-9(ok2853)*.

### Male mating behavior assays

Male mating behavior assays were performed as described previously (Hodgkin, 1983) with minor modifications. To prepare a small bacterial lawn for mating, NGM plates were seeded with 10 µL of fresh OP50 liquid culture and allowed to dry. 10 1-day-old adult *unc-31(e169)* hermaphrodites were added to the lawn. After a 30-minute acclimation period, a 1-day-old adult male was placed on the lawn and observed for 5 minutes. Contact response was recorded when the male encountered a hermaphrodite and transitioned from forward movement to scanning the hermaphrodite. Location of vulva was recorded when males successfully located and stopped scanning when they reached the vulva. For each genotype, 4 replicates of 10 males each were assayed with the experimenter blinded to the genotype.

### Live imaging

One day prior to imaging, L4 animals were isolated onto NGM plates seeded with OP50. On the day of imaging, adult animals were placed into a 1 uL droplet of 10 mM levamisole on a coverslip and then mounted on a 10% agarose pad. Imaging was performed with a Zeiss LSM 880 microscope with Airyscan Super-Resolution Detector with 7 Single Photon Lasers (405, 458, 488, 514, 561, 594, 633 nm), Axio Observer 7 Motorized Inverted Microscope, Motorized X-Y Stage with Z-Piezo, T-PMT. Images were acquired using a 63x/1.4 Oil Plan-Apochromat objective. Raw image files were processed with ZenBlack 2.0 software (Airyscan Processing). Processed images were imported into ImageJ for analysis and to adjust display for presentation. Images are displayed as maximum intensity projections of Z-stacks unless otherwise noted in the figure legend.

### CIL-7::GFP live imaging

Animals were prepared for imaging in the same manner described under “Live Imaging.” For the imaging of CIL-7::GFP, epifluorescence images were acquired using a Zeiss Axioplan2 microscope with 63x oil-immersion objective and Photometrics Cascade 512B CCD camera. Images are displayed as maximum intensity projections of Z stacks.

### PKD-2::GFP quantification

PKD-2::GFP fluorescence intensity was quantified using Fiji (Schindelin et al., 2012). For each image, a maximum intensity projection of the Z-stack was generated. Then, the area containing the cilium was duplicated and the threshold function was used to define an ROI. The area and mean intensity of the ROI were measured, and an equivalent ROI from a region of the image that didn’t contain any worm was measured as background. The total cilium and background fluorescence were calculated as area*mean intensity for each ROI. Then, the background intensity was subtracted from the cilium intensity.

### TSP-6::mNG phasmid cilia length measurement

Phasmid cilia length as visualized by TSP-6::mNG fluorescence was measured using ImageJ. For each image, a maximum intensity projection of the Z-stack was generated. Length measurements were performed using line scans across cilia and the full width at half maximum of the line scan from cilium tip to periciliary membrane is presented.

### Extracellular vesicle imaging and quantification

Extracellular vesicle imaging and quantification were performed as described previously (Wang et al., 2024b). Animals were prepared for imaging in the same manner described under “Live Imaging” and imaged at the initial 0 hour timepoint and 1 hour later. EV quantification was carried out using ImageJ. For each image, a maximum intensity projection of the Z-stack was generated. The region containing EVs was selected for analysis and the threshold function applied such that each EV appears as an individual ROI (the same threshold was used across genotypes and the 0 h and 1 h timepoints). The total number of ROIs was recorded as the number of EVs per animal.

### Transmission Electron Microscopy

*mec-9(ok2853)* and wild-type young adult males were fixed using high-pressure freeze fixation and freeze substitution in 2% OsO4 + 2% water in acetone as the primary fixative (Weimer, 2006). Samples were slowly freeze-substituted in an RMC freeze substitution device, before infiltration with Embed-812 plastic resin. For TEM, serial sections (70-75 nm thickness) of fixed animals were collected on copper slot grids coated with formvar and evaporated carbon and stained with 4% uranyl acetate in 70% methanol, followed by washing and incubating with aqueous lead citrate. Images were captured on a Philips CM10 transmission electron microscope at 80kV with a Morada 11-megapixel TEM CCD camera driven by iTEM software (Olympus Soft Imaging Solutions). Images were analyzed using ImageJ and prepared for publication with Adobe Illustrator.

### Statistical analysis

R and RStudio were used to carry out statistical analyses. For PKD-2::GFP quantification in rays (Figure 1E-G, Figure S2E-G), when the two-way ANOVA indicated a significant effect of ray identity (R3B v. R4B v. R5B), each ray is plotted separately.

## Supporting information

source data for figures

## Acknowledgements

For outstanding technical support, we thank Gloria Androwski. For assistance with male mating assays, we thank Elizabeth desRanleau. For assistance with live microscopy, we thank Juan Wang. For providing strains, we thank: Chalfie Lab (Columbia University), Heiman Lab (Harvard Medical School/Boston Children’s Hospital), Caenorhabditis Genetics Center (CGC), *C. elegans* Gene Knockout Project at the Oklahoma Medical Research Foundation/International *C. elegans* Gene Knockout Consortium (Consortium, 2012). We thank all Barr lab members for constructive discussions and comments on the manuscript. We used key resources WormBase (Sternberg et al., 2024) and WormAtlas (Altun) throughout the project.

---

**Figure S1.**
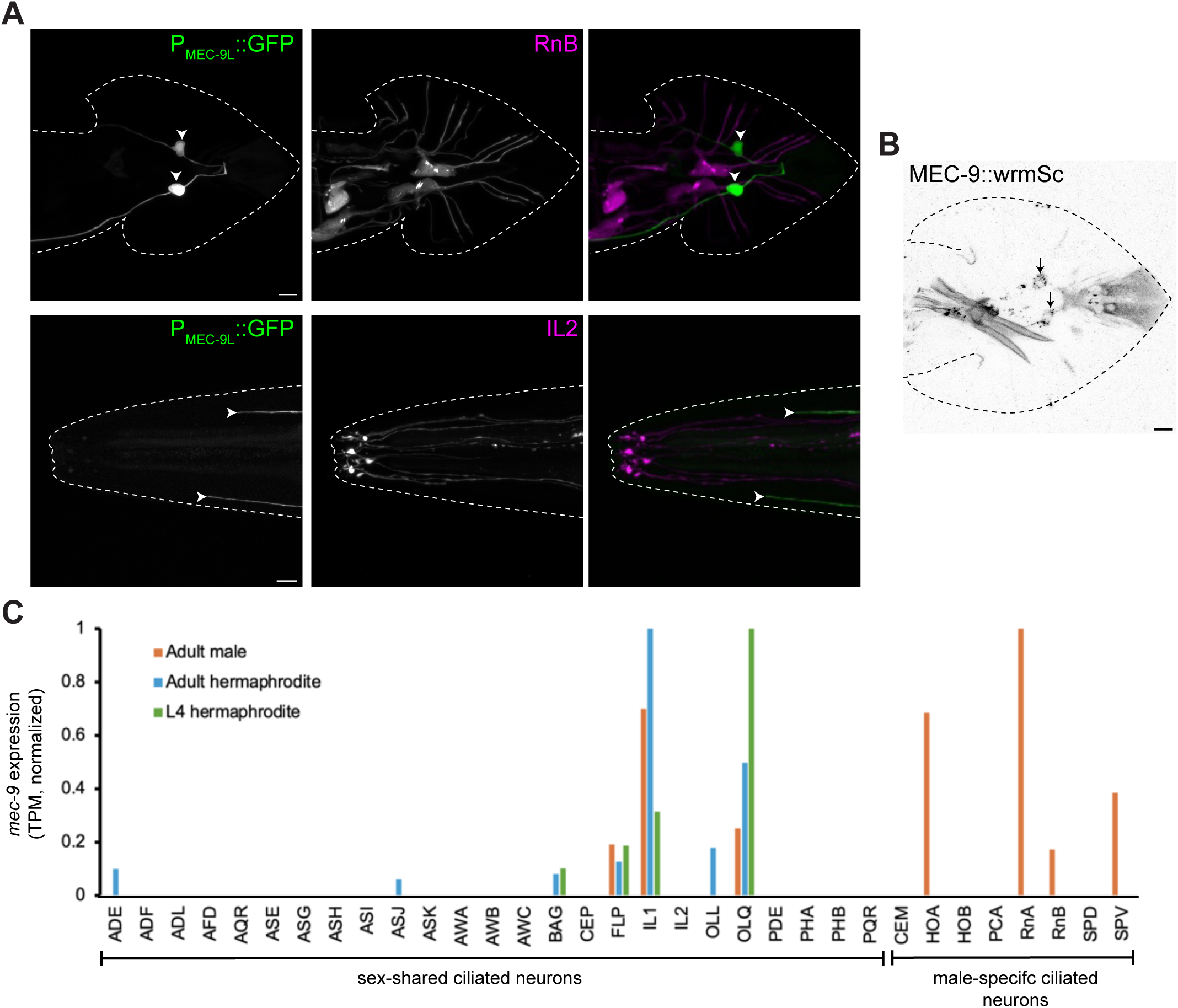
(A) The long isoform encoded by *mec-9* is expressed in touch receptor neurons. (Top) Confocal image of MEC-9L transcriptional reporter (green) in adult male tail, co-expressed with *P_klp-6_::tdTomato* (magenta) which labels RnB neurons (PT4377). Arrowheads: PLM. Scale bar = 5 µm. (Bottom) Confocal image of MEC-9L transcriptional reporter (green) in adult male head, co-expressed with *P_klp-6_::tdTomato* (magenta) which labels IL2 neurons (PT4377). Arrowheads: ALM processes. Scale bar = 5 µm. (B) Confocal image of endogenously-tagged MEC-9::wrmScarlet in adult male tail (PT4128). Arrows: PLM. Scale bar = 5 µm. (C) *mec-9* expression in ciliated sensory neurons in transcripts per million (TPM), values normalized to maximum. Data from the CeNGEN adult male and hermaphrodite data sets (Olson) and L4 hermaphrodite data set (Taylor et al., 2021).

**Figure S2.**
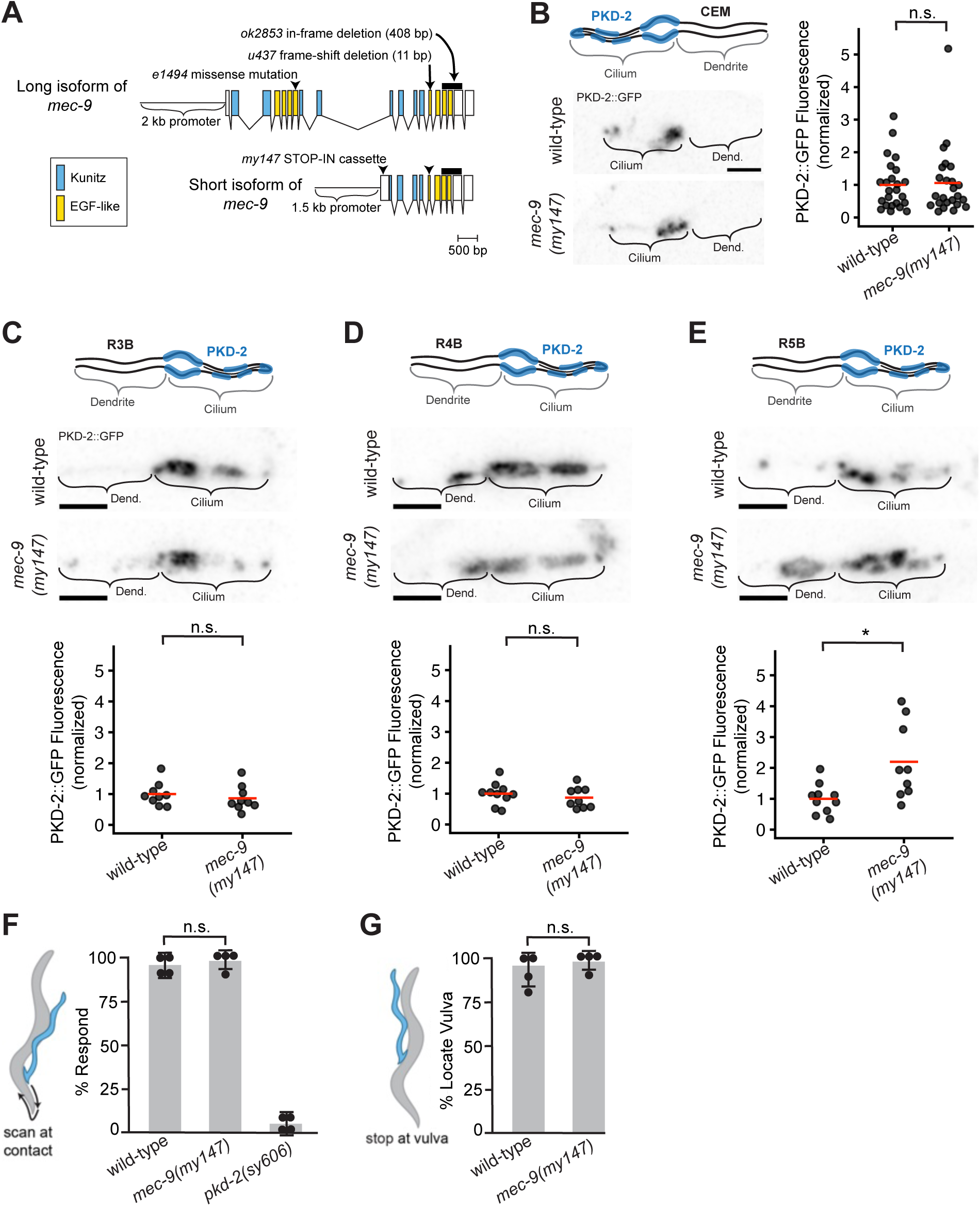
A STOP-IN allele targeting the short isoform encoded by *mec-9* affects PKD-2::GFP localization only in R5B cilia and does not cause defects in male mating behaviors. (A) Intron-exon schematic of isoforms encoded by *mec-9* with alleles indicated. (B) Confocal images and quantification of PKD-2::GFP at CEM cilia in wild-type (PT3168) and *mec-9(my147)* mutants (PT4125). N ≥ 22 cilia. One-way ANOVA. The wild-type data is the same as presented in Figure 1. (C, D, E) Confocal images and quantification of PKD-2::GFP in R3B cilia (C), R4B cilia (D), and R5B cilia (E) in wild-type and *mec-9(my147)* mutants. N ≥ 9 cilia. Two-way ANOVA revealed no main effect of genotype but a significant interaction between genotype and ray identity. Post-hoc testing (TukeyHSD) revealed a significant difference in R5B (p = 0.003). For B-E, fluorescence values are normalized to the wild-type mean and red bars in dot plots indicate the mean. Scale bars = 2 µm. (F and G) *mec-9(my147)* mutants (PT4125) are not defective in response to mating partner (F) and location of vulva (G) when compared to wild-type controls (PT3168). *pkd-2(sy606)* (PT9) is a negative control for mating assays. N = 4 mating assays, 10 animals per assay. One-way ANOVA and post-hoc testing with TukeyHSD.

**Figure S3.**
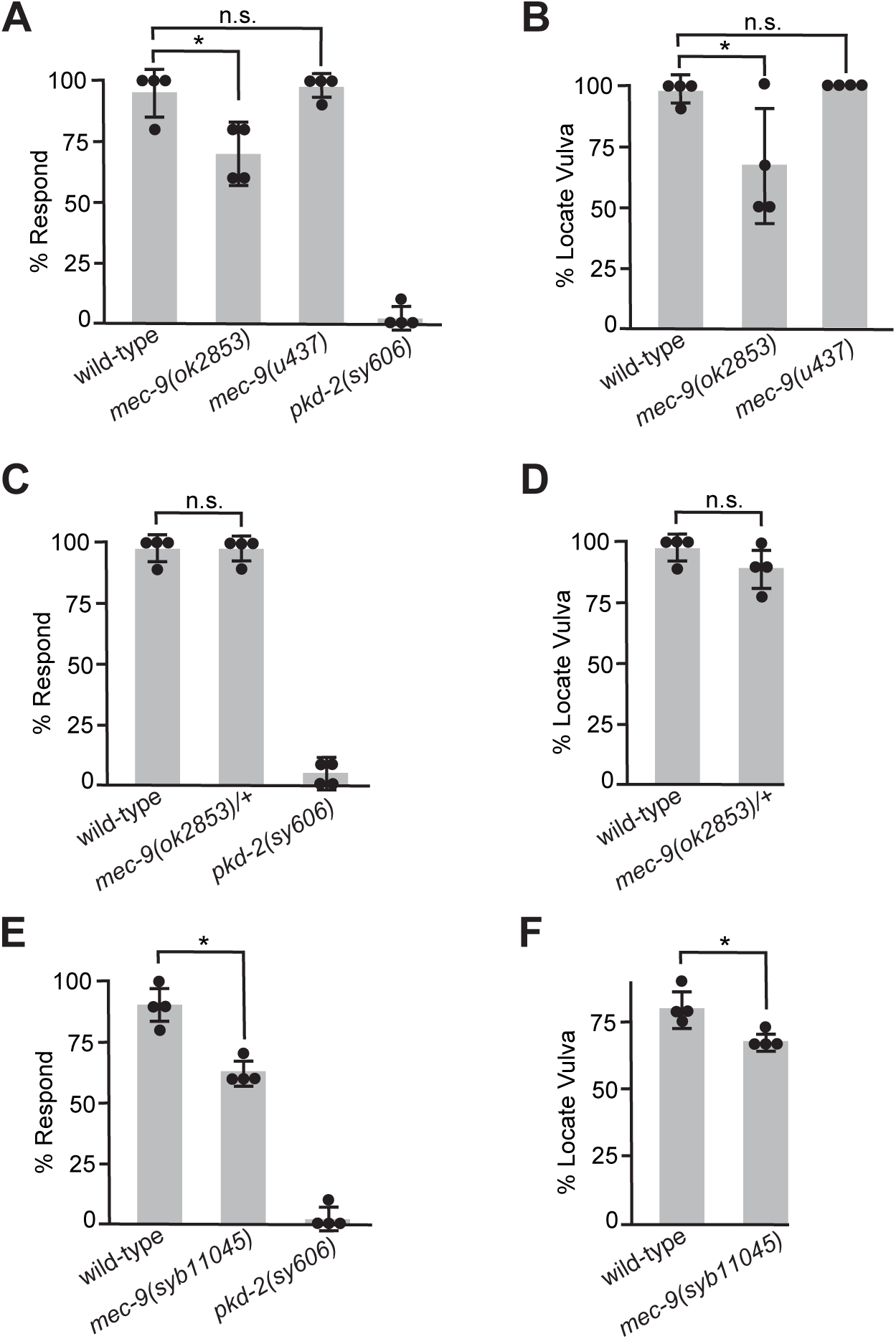
(A-B) *mec-9(u437)* (PT4241) mutants do not exhibit defects in contact response to mating partner (A) or location of vulva (B). Wild-type, *mec-9(ok2853)* and *pkd-2(sy606)* data is the same as in Figure 1. (C-D) *mec-9(ok2853)* is recessive. Males heterozygous for the *ok2853* mutation do not exhibit defects in contact response or location of vulva. To generate males heterozygous for *mec-9(ok2853)*, mutant hermaphrodites (PT4164) were crossed with wild-type males carrying *P_unc-122_::GFP* which labels the coelomocytes (PT3168). Cross progeny expressing *P_unc-122_::GFP* were selected for mating assays. (E-F) *mec-9(syb11045)* (PT4447) mutants exhibit defects in contact response and vulva location. *mec-9(syb11045)* has the same 408 base-pair deletion as *mec-9(ok2853)*. For all mating assays, N = 4 assays, 10 animals per assay. One-way ANOVA and post-hoc testing with TukeyHSD, * = p ≤ 0.01.

**Figure S4.**
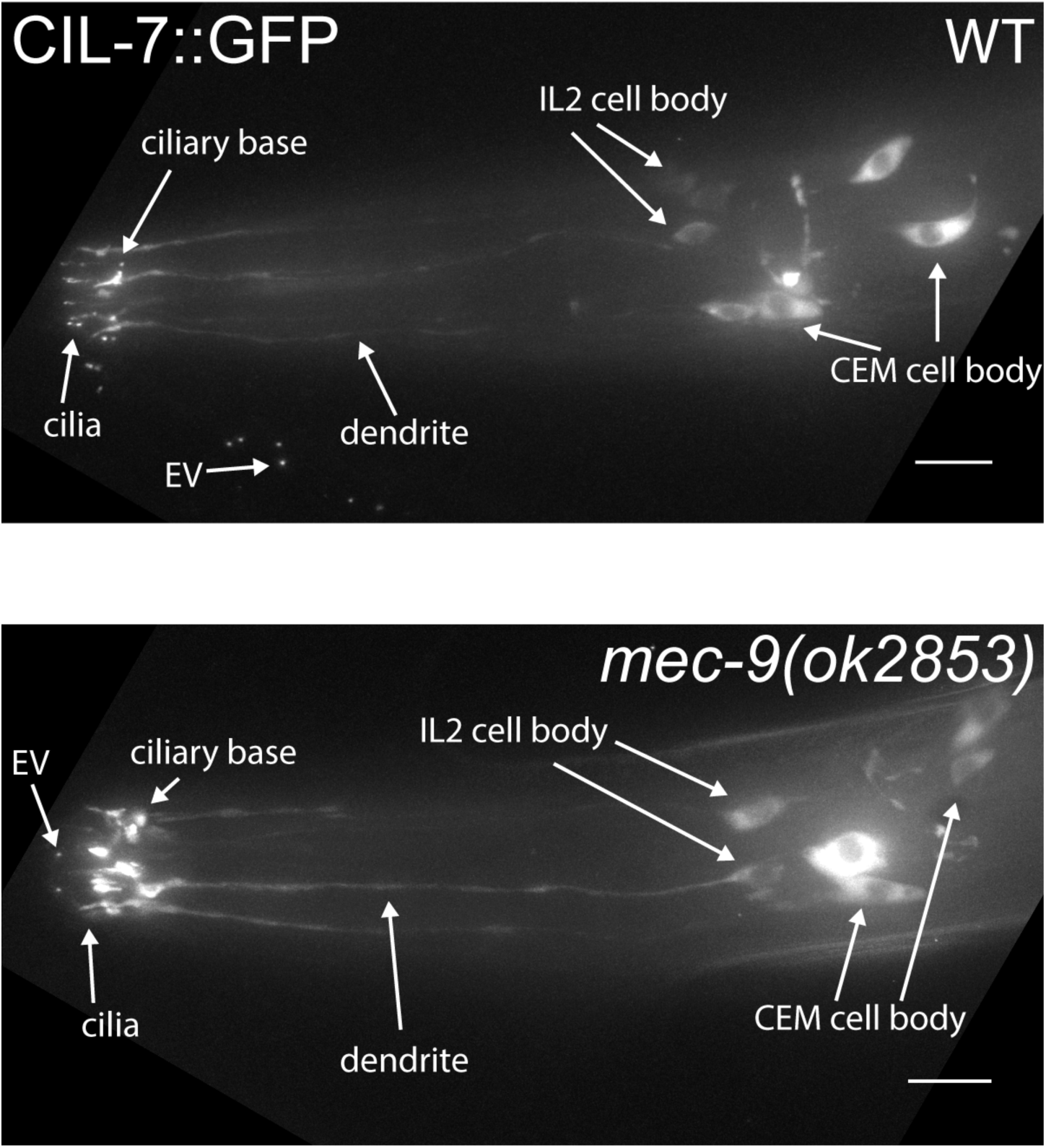
*mec-9(ok2853)* impacts CIL-7::GFP localization in IL2 cilia. *mec-9(ok2853)* mutant (PT3213) displays abberrant accumulation of CIL-7::GFP in IL2 and CEM cilia compared to wild-type control (PT2679). Scale bar = 10 µm.

**Figure S5.**
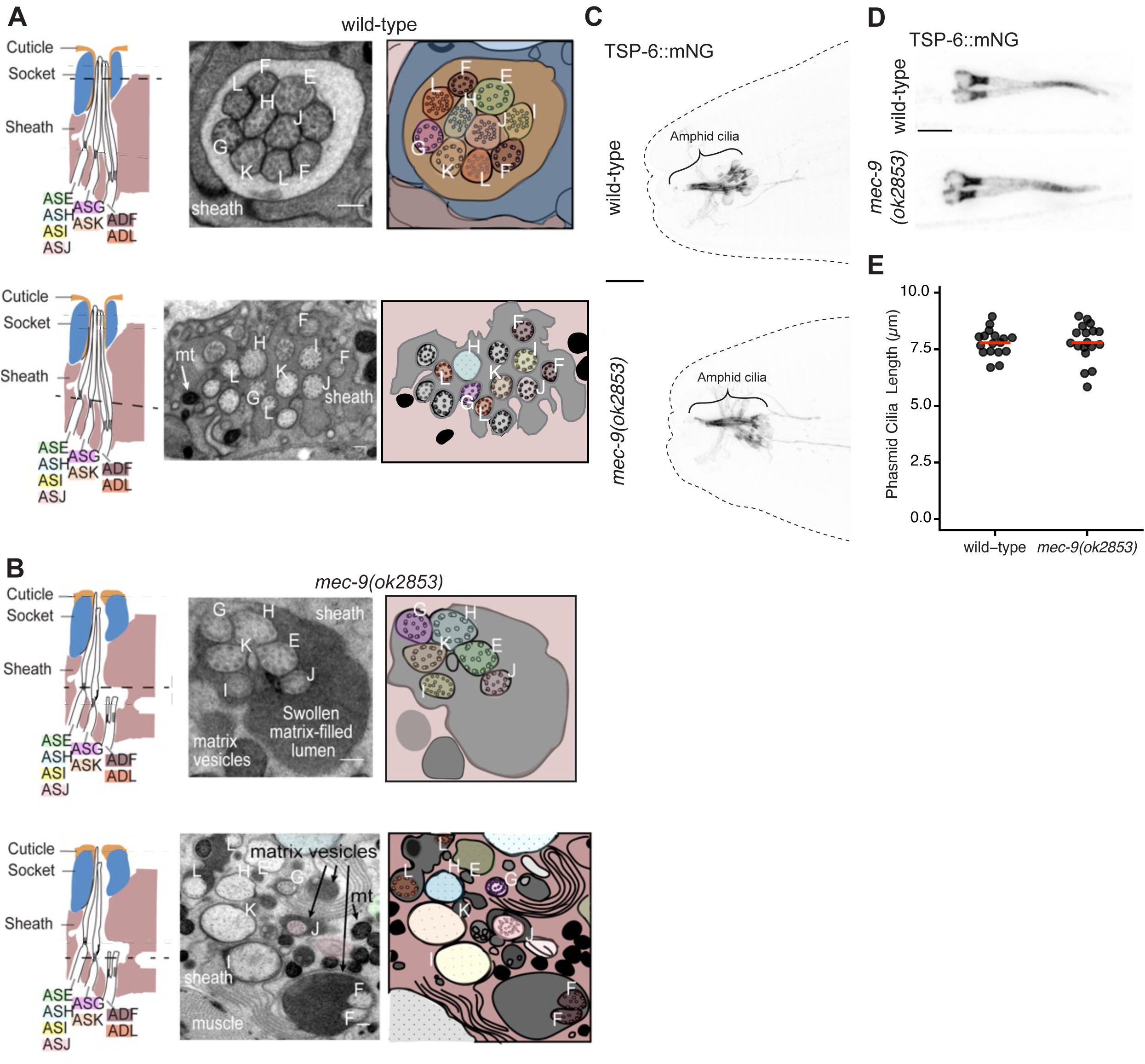
*mec-9(ok2853)* impacts amphid sensory organ morphology but does not affect TSP-6::mNG localization to amphid cilia or phasmid cilia length. (A-B) TEM cross sections of the amphid sensory organ at the level indicated by the dashed line in the cartoon. (A) Top: Wild-type middle segment contained 10 cilia with microtubules arranged concentrically. Bottom: Wild-type transition zone (TZ) region shows electron-dense matrix material surrounding the amphid cilia. (B) Top: *mec-9(ok2853)* mutant middle segment: six out of ten cilia were visible in the lumen and the lumen was swollen. Bottom: *mec-9(ok2853)* TZ and distal dendrite region: the shortened ADF and ADL cilia are visible. For A and B, N = 4-5 sense organs examined across 2-3 animals per genotype. Scale bars = 200 nm. (C) Confocal images of TSP-6::mNG in amphid cilia in wild-type (PT3914) and *mec-9(ok2853)* mutants (PT4189). Scale bar = 5 µm. (D) Confocal images of TSP-6::mNG in the phasmid cilia of wild-type and *mec-9(ok2853)* mutants. Scale bar = 2 µm. (E) Quantification of phasmid cilia length in wild-type and *mec-9* mutants using TSP-6::mNG as a marker (see Methods). N ≥ 17 cilia per genotype. T-test = NS. Red bars indicate the mean.

